# Garlic (Allium sativum L.) yield and quality as affected by different integrated fertilizer levels at Wachemo university, southern Ethiopia

**DOI:** 10.1101/2024.10.05.616817

**Authors:** S. Abrham, J. Mihret, B. Abraham

**Affiliations:** Horticulture Department, Agriculture College, Wolaita Sodo University, Ethiopia P.O. Box 138, Wolaita Sodo, Ethiopia; Plant Sciences Department, Agriculture College, Wachemo University, Ethiopia

**Keywords:** Bulb yield, chicken manure, fertilizer, quality, production

## Abstract

Production and productivity of garlic was constrained by many factors, among which lack of integrated nutrient recommendation is the most important in Ethiopia. Thus, on site trial was executed at Wachemo University Lembuda demonstration field during 2021/2022 with the aims to identify the effects of NPS fertilizer and chicken manure levels on growth, bulb yield and quality; to determine the economic optimum combination of NPS and chicken manure rate for garlic productivity in the target location. The treatments included four levels of chicken manure (0, 5, 10 and 15 t/ha) and five levels of NPS fertilizer (0, 50,100, 150 and 200 kg/ha) arranged factorially using a randomized complete block design (RCBD) and replicated three times. Growth, bulb yield and quality data were collected and subjected to analysis using SAS version 9.4. With the exception of days to maturity, leaf number and non-marketable bulb yield in the case of chicken manure, the majority of the investigated parameters were significantly (P < 0.05) impacted by the main effects of chicken manure and NPS fertilizer. Exception marketable and total bulb yield, the interaction impact was not significant for nearly all of the examined variables. On the other hand, the bulb yield was considerably raised by raising the rate of chicken manure in combination with NPS fertilizer. As a result, combined application of 15 t/ha chicken manure and 150 kg/ha NPS fertilizer produced the maximum marketable bulb (19.52 t/ha) and total yield (20.27 t/ha), followed by marketable yield (18.53 t/ha) and total yield (19.55 t/ha) obtained from plants treated with 200 kg/ha of NPS fertilizer and 15 t/ha of chicken manure, with a 927 % yield advantage over plants receiving no fertilizer. On the other hand, the control treatment produced the lowest total yield (3.89 t/ha) and marketable yield (1.90 t/ha). Plots with 15 t/ha chicken manure had the highest bulb dry matter (35.31%) and TSS (22.47° Brix). In terms of economic analysis, the plants that got 15 t/ha of chicken manure and 150 kg/ha of NPS fertilizer had the highest net benefit (524224.8 ETB/ha) with an MRR of 10,159.1%. Hence, this combination was found to be the best and cost-effective one for smallholder farmers in the study area that could be recommended to boost garlic production and productivity.

## Introduction

After onions, garlic (*Allium sativum* L.) is the second most popular bulb crop in the Alliaceae family [1]. Ethiopia is among the many countries in the globe where it is extensively grown. Garlic was estimated to be grown on 1546741 hectares worldwide, with a total annual production of 28494130 tons [2]. In Ethiopia, 32,080.40 ha of land was under garlic cultivation with a production of about 196,416.15 tonnes resulting in an average productivity of 6.12 t/ha, which is far lower than the global average yield of 18.4 t /ha [3].

All parts of this plant (cloves, leaves, stem and seed) have been used and consumed either fresh or cooked for health benefits for diseases like diabetes, cancer, and ulcer rheumatism [4]. Alemu *et al*. [5] reported that garlic is cultivated for home consumption as a spice and one of the major bulb crops containing various minerals, alkaloids, tannins, carotenoids, saponin, flavonoids, steroids, and cardenolides [6].

One of the factors limiting the productivity of various crops in Ethiopia is low soil fertility, which is primarily caused by crop residue removal from farmland, crop nutrients being removed from the soil by crops, erosion of surface soil, and the absence of a crop rotation system [7, 8]. Ethiopian farmers have a tradition of adopting crop rotation, intercropping, adding farmyard manure, other organic fertilizers, and fallowing to maintain or improve the fertility of the soil in order to reduce these issues [9].

Integrated nutrient application has complementarity effects that ensure the balanced provision of nutrients essential to plants with effective reactions in advancing mineral use efficiency of crops and improve soil fertility and health [10, 11]. Some attempts done so far in garlic crop production around the world indicated that the integrated use of nutrients have a valuable impact on bulb yield and quality. The integrated use of vermicompost and NP fertilizers was investigated, and a significant yield advantage was observed [10]. Again, Getachew and Asefa [11] have reported the advantages of the combined addition of NPS mineral fertilizers and farm yard manure for garlic production and productivity in Ethiopia. Furthermore, Sitaula [12] reported the agronomic and economic advantage of the mixed application of poultry manure and urea on the growth, bulb yield and quality of garlic in Nepal.

Garlic crop yield appears to be directly increased by chicken manure either by speeding up the metabolic process, by increasing cell permeability and hormone growth action or combination of all three activities [13]. Additionally, it provides several nutrients and enhances the physical characteristics of soil, including soil aggregation, permeability, and water-holding ability [14].

However, garlic cultivation in the country is constrained by many challenges, among which the lack of integrated nutrient management practices is one of the most limiting factors in determining its production and productivity. Small-scale farmers use poor agronomic practices with inorganic fertilizers or compost for the production of garlic, which contributed to low productivity and quality in the current study areas (personal observation). Furthermore, no research has been conducted in the area regarding the amount and application of chicken manure, mineral nutrients, and their combination. Currently, the ever increasing cost of inorganic fertilizers in Ethiopia, in general, calls for research for alternative options to replace them by organic fertilizers, among which chicken manures are the most important ones production of the crop in the experimental site. Thus, in order to increase crop yield and quality, a systematic study of chicken manure and inorganic fertilizers (NPS) is essential. Therefore, the current study was aimed to determine how varied ratios of chicken manure and NPS fertilizer together affect garlic bulb yield and quality. It also aimed to investigate the influence of chicken manure and levels of NPS on bulb yield and quality, and to determine the economic optimum of chicken manure and NPS level for garlic cultivation.

### Materials and methodology

#### Description of the study site

The field experiment was executed at Lembuda Agriculture field practical nursery site of Wachemo University in Lemo woreda, Hadiya zone, Southern Ethiopia, in 2021/2022 in two seasons. Its altitude is 2381 m.a.s.l. and is located at 48^0^ 5’2.094’’ E longitude and 3^0^20’45.16’’ N latitude. The recorded annual mean temperature ranges from 15 – 18 ^0^c and average annual rainfall was 1150 mm (Ethiopian National Meteorology Agency, 2018). The type of experimental soil was clay loam in texture having a pH of 5.98.

**Figure 1:**
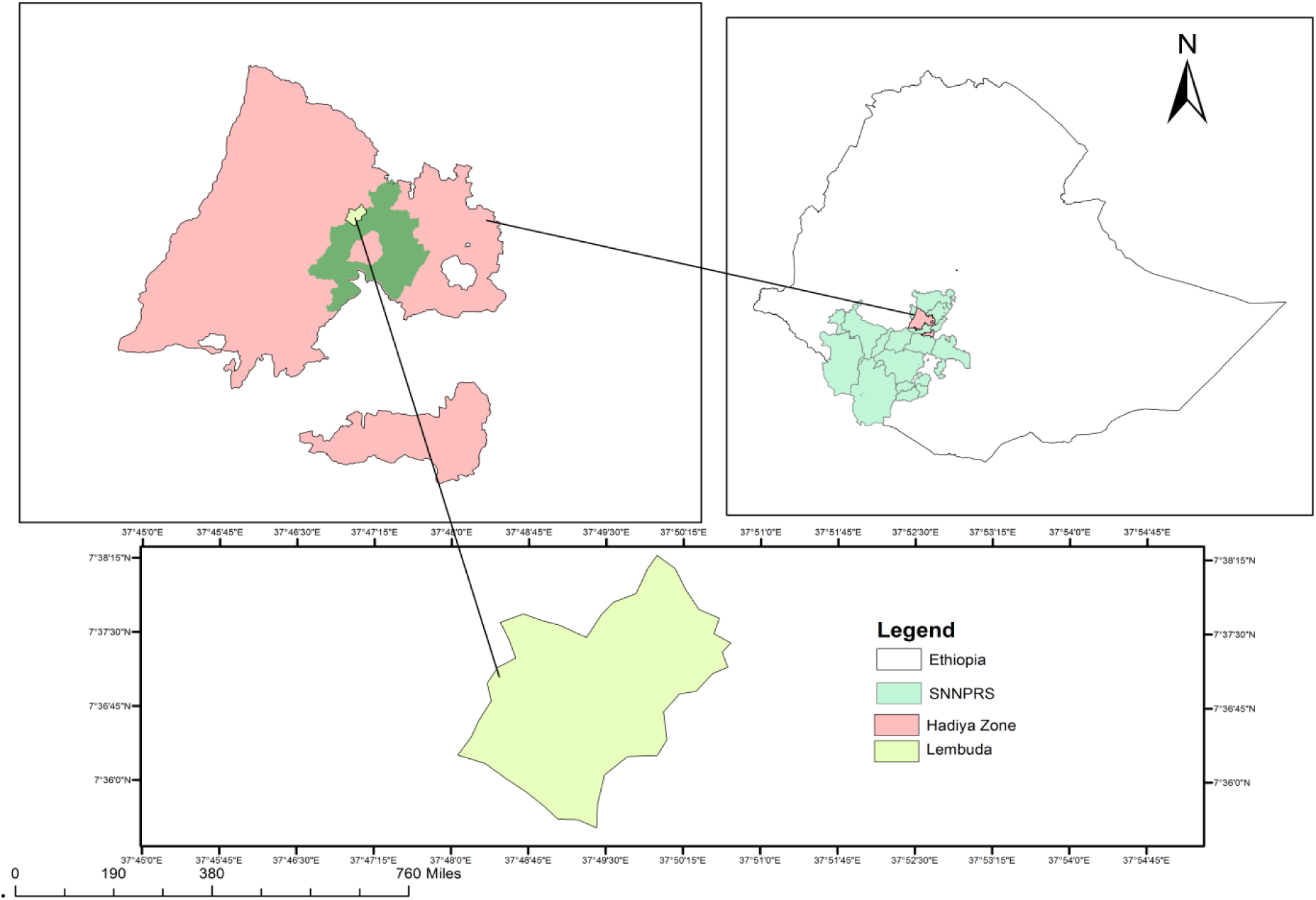
Map of study area

#### Experimental design and treatments

The experiment had two major factors. The first factor consisted of five different rates of NPS (0, 50, 100, 150 and 200 kg/ha) and the second had four rates of decomposed chicken manure (0, 5, 10 and 15 ton/ha) which resulted in 20 treatments in total. The trial was set up in a factorial arrangement in a randomized complete block design (RCBD) having three replications. The spacing used during planting was 10 cm between plants and 30 cm between rows. The number of rows per plot was five and the number of plants per row was 15 giving a total of 75 plants per plot. The gross plot size was 1.5 m × 1.5 m (2.25 m^2^). The space between blocks and plots was 0.75 and 0.5 m, respectively. The total number of plots was 60 and total experimental area was 240 m^2^. The NPS fertilizer rates required for the treatments were applied during planting. All cultural practices including weeding, harrowing and watering were consistently applied for all plots as recommended by Debre Zeit Agricultural Research Centre [15].

### Data measurements

#### Phenological and growth parameters

**Days to physiological maturity:** this parameter represented the number of days needed from planting until 90% of the plants in each plot displayed leaf senescence.

**Plant height (cm):** using a ruler, the height of ten randomly chosen plants from each plot’s net plot area was measured at maturity from the soil’s surface to the tip of the plant.

**Number of leaves per plant:** the average number of leaves per plant was calculated by counting the leaves on ten mature plants.

**Leaf length (cm):** from ten randomly selected plants, the average leaf length at physiological maturity was measured in centimeters.

#### Bulb yield, yield components and quality

**Bulb length (cm):** using a caliper, the length of ten bulbs was measured from the base to the beginning of the neck.

**Bulb diameter (cm):** using a caliper, the diameter of ten randomly selected bulbs was measured at the widest point in the middle of the bulb.

**Clove number per bulb:** the average clove number per bulb was calculated by counting all of the cloves that were produced from ten randomly selected plants.

**Clove weight (g):** the weight of each of the cloves taken from ten bulbs was measured and the average weight was recorded.

**Marketable bulb yield (t/ha):** bulbs were thinly scattered for drying and cured for 10 days under ambient conditions after being collected at 90% physiological maturity of the plants in the net plot area. Then bulbs with a larger size that had not been damaged chosen, weighed and translated into t/ha.

**Unmarketable bulb yield (t/ha):** this refers to the bulbs that were small-sized, imperfect, mechanically damaged, or attacked by pests.

**Total bulb yield (t/ha):** the total amount of marketable and unmarketable yield from the net plot area converted in to the hectare basis to determine the total bulb yield.

**Bulb dry matter content (%):** is the average bulb dry matter weight of ten randomly selected plants, the average mature bulb weight of which was measured after the bulbs were dried in an oven with forced hot air circulation for seventy-two hours at 70 ^0^C for ten days to dry the bulbs.

TSS (°Brix): was obtained by taking readings on a digital refractometer that had automatic temperature correction, straight from the homogenized garlic juice. As per the AOAC (2002) approach, the outcomes were represented as (°Brix).

#### Data analysis ANOVA

The data gathered and summarized were employed to analysis of variance (ANOVA) using Statistical Analysis Software SAS, version 9.4; mean separation was executed by using LSD-test at P < 0.05 probability.

#### Economic analysis

Garlic yield and efficiency can also be guaranteed by performing a cost-benefit analysis on the application of NPS inorganic fertilizers and chicken manure, following the guidelines provided by CIMMYT [16]. The study’s primary inputs were labor, chicken manure, urea, NPS fertilizer, and garlic cloves. The costs of these materials were calculated at Ethiopian Birr local market prices and totaled to determine the study’s total variable cost. The marketable bulb was modified by 10% in order to account for the true productivity of farmers as reported by CIMMYT [16], given the clear differences in field management between research stations and farmers. The average adjusted marketable bulb (t ha−1) was multiplied by the local farm gate price of Ethiopian Birr to estimate the gross return per hectare basis. Therefore, the gross return was subtracted from the total variable cost to determine the net benefit.

The percentile ratio of the change in net benefit (NB) to total variable cost (TVC) was used to estimate the marginal rate of return (MRR).

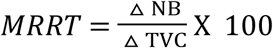

## Result and discussion

### ANOVA results

Generally, the investigation result indicated the main effect of decomposed chicken manure significantly (P < 0.05) influenced all traits except days to maturity, number of leaf plant^-1^ and unmarketable bulb yield. In contrast, the influence of NPs fertilizer rates had significant (P < 0.05) effect on all the investigated parameters. The results also showed that, with the exception of marketable and total bulb yield, the interaction of fertilizer rates and chicken manure was not significant for the majority of the parameters (Table 1).

**Table 1:**
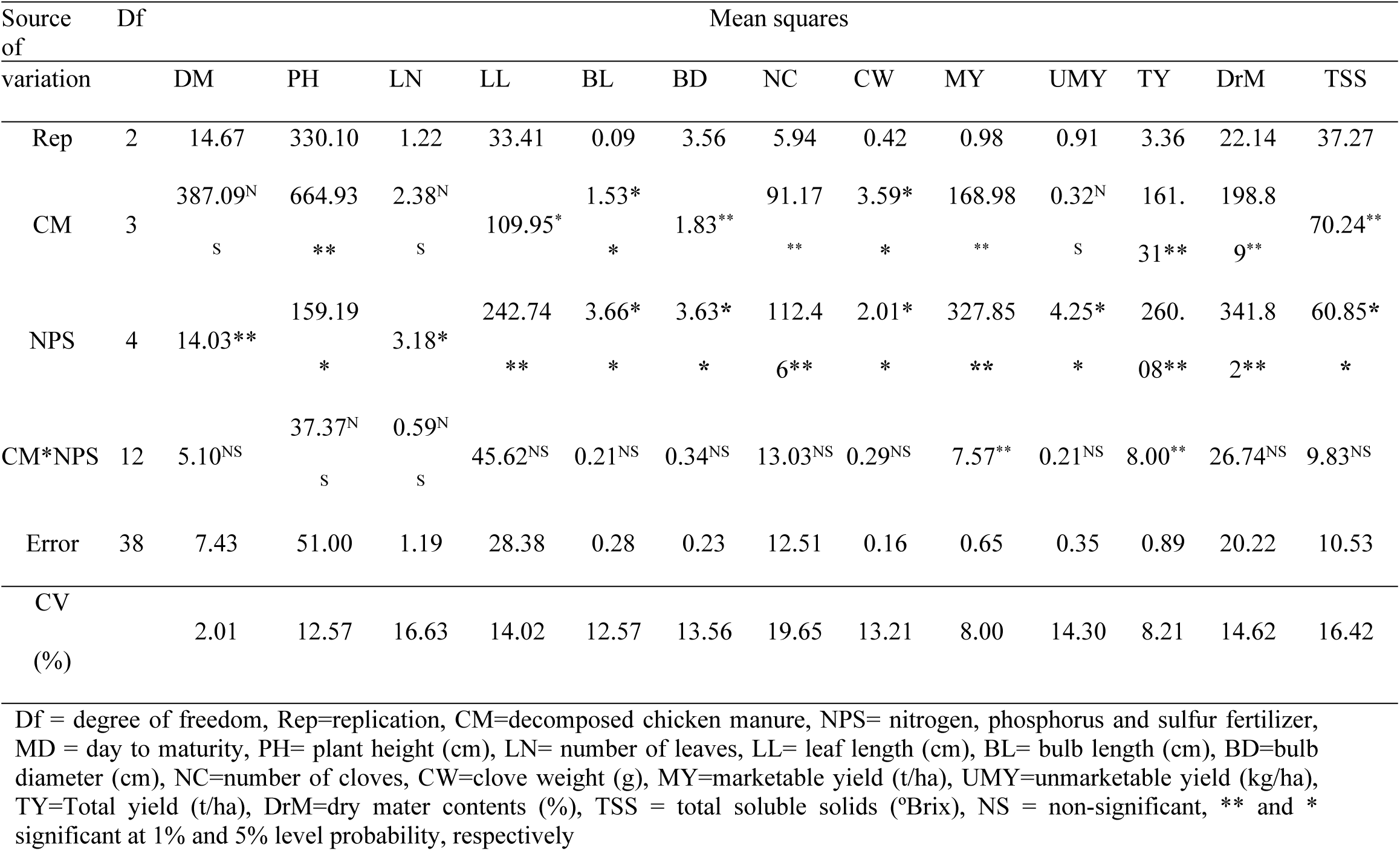
ANOVA result indicating the mean values of 12 parameters considered during the experiment in 2021/2022.

### Crop phenology and growth

#### Plant height (cm)

The results of the analysis of variance indicated that the main impacts of NPS fertilizer rates and chicken manure had a significant (P < 0.01) impact on plant height. However, their interaction had no significant impact on plant height (Table 1). The application of 15 t/ha chicken manure resulted in the highest mean plant height (62.77 cm), which was statistically comparable to 10 t/ha chicken manure (60.59 cm). The shortest plant height, with a mean value of 47.71 cm was obtained by zero chicken manure (Table 2).

**Table 2:** The influence of NPS fertilizers and chicken manure rates on the days-to-maturity (DM), plant height (PH), number of leaf (LN), leaf length (LL) in 2021–2022.

| Treatments | MD | PH(cm) | LN | LL(cm) |
| --- | --- | --- | --- | --- |
| Chicken manure rate (t/ha) |  |  |  |  |
| 0 | 141.20<sup>a</sup> | 47.71<sup>c</sup> | 6.05 | 35.05<sup>b</sup> |
| 5 | 137.33<sup>b</sup> | 56.15<sup>b</sup> | 6.64 | 36.75<sup>b</sup> |
| 10 | 133.60<sup>c</sup> | 60.59<sup>a</sup> | 6.59 | 38.95<sup>ab</sup> |
| 15 | 129.33<sup>d</sup> | 62.77<sup>a</sup> | 7.01 | 41.29<sup>a</sup> |
| LSD<sub>(0.05)</sub> | 2.02 | 5.27 | NS | 3.94 |
| NPS rate (kg/ha) |  |  |  |  |
| 0 | 136.67 | 52.1<sup>c</sup> | 6.03<sup>b</sup> | 31.23<sup>c</sup> |
| 50 | 136.25 | 55.22<sup>bc</sup> | 6.28<sup>b</sup> | 36.64<sup>b</sup> |
| 100 | 135.25 | 56.40<sup>abc</sup> | 6.38<sup>b</sup> | 38.02<sup>ab</sup> |
| 150 | 134.5 | 58.45<sup>ab</sup> | 7.34<sup>a</sup> | 41.81<sup>a</sup> |
| 200 | 134.17 | 61.86<sup>a</sup> | 6.82<sup>ab</sup> | 42.33<sup>a</sup> |
| LSD<sub>(0.05)</sub> | NS | 5.9 | 0.9 | 4.4 |
| CV | 2.01 | 12.57 | 6.63 | 14.02 |

Values in the same column with the same letters are statistically similar at 5% probability level, CV (%) = Coefficient of variation, LSD<sub>(0.05)</sub> = Least Significant Difference.

This could be explained by the fact that chicken manure includes various mineral nutrients, carbon, and nitrogen, all of which may have helped to encourage plant growth. The results of Somen and Hitesh [17], who observed that poultry manure delivered all needed nutrients resulting in an increase in plant height, were consistent with this outcome. Similarly, the application of 15 t/ha of cattle dung resulted in the highest mean height of garlic (67 cm) [18]. The final result was in line with the research conducted by Adewale *et al*. [19], who found that a treatment with 20 t/ha of poultry manure resulted in a height of 79.8 cm in garlic plant.

Garlic plant height was significantly (P < 0.05) influenced by the rates of NPS fertilizers (Table 1). The treatment that received 200 kg/ha had the tallest plants (61.89 cm), while the treatment that received zero or control fertilizer had the least plant height (52.1 cm) (Table 2). This outcome might be the result of indirect effects of phosphorus and sulfur on crop growth and development, as well as the presence of nitrogen, which promotes vegetative growth. Similar to the current study, Indupulapati [20] showed that potassium, phosphorus, sulfur, and nitrogen are the main nutrients needed by plants and are crucial building blocks of proteins, enzymes, carbohydrates, and vitamins. The present investigation aligned with the findings of Fikre *et al.* [21], which revealed that the maximum plant height was achieved by applying 130 kg/ha of NPS fertilizer, and in another study, the highest plant height (65.7 cm) was achieved by applying 187.5 kg/ha of NPSB fertilizer on onions. Furthermore, <u>Shege</u> *et al.* [47] in their finding reported that the highest plant height (68.5 cm) was achieved by the higher doses of NPS (140:122.6:22.6) combinations in garlic plant.

#### Leaf number per plant

The number of leaves per plant was not significantly affected by the main effects of chicken manure, but it was significantly affected (P < 0.05) by the effects of NPS rates (Table 1). As a result, the application of 150 kg/ha NPS produced the highest number of leaves per plant (7.34), which is statistically comparable to the application of 200 kg/ha (6.82). The application of zero fertilizer produced the lowest number of leaves per plant (6.03), which was statistically non-significant when compared with 50 and 100 kg/ha NPS fertilizer rates (Table 2). The observed variability in the number of leaves per plant could be related to the crop’s response to nitrogen, phosphorus, and sulfur fertilizers. As a part of chlorophyll, nitrogen helps produce photosynthesis, which promotes stronger vegetative growth. Because it is a necessary component of both cellular protein and nucleic acid, phosphorus directly affects meristematic activity, which increases the number of leaves per plant. Consistent with our study, Getachew and Temesgen [22] found that applying 78.75-69-12.75 kg N-P-S ha^−1^ resulted in the maximum leaf number per plant (7.03), whereas the control plot, which received no fertilizer, had the lowest leaf number (6.0).

#### Leaf length (cm)

The chicken manure and NPS fertilizer rates significantly (P < 0.01) influenced leaf length in their main effects. However, the treatment interaction did not affect leaf length significantly (Table 1).

In the case of chicken manure, the longest mean leaf length (41.29 cm) was revealed by the treatment of 15 t/ha and that of 10 t/ha recorded 38.95 cm, which was statically on par. Whereas the lowest leaf length with a mean value (35.05 cm) was recorded from zero chicken manure again, followed by 36.75 cm recorded by application of 5 t/ha which was statistically non-significant (Table 2). This might be the result of added manure that increased plant vegetative growth due to the availability of nutrients released, Frank [23]. This is consistent with the results of Somen and Hitesh [17], who found that 20 t/ha of vermicompost applied gave the longest leaves (55.6 cm) whereas, the shortest leaves (43.7 cm) was obtained by zero or control treatment. According to Michael [24], the plants from the 20 t/ha poultry manure treatments had leaves that were noticeably longer (44.3 cm) than those from the other treatment.

The NPS at 200 kg/ha had numerically greatest leaf length (42.33 cm) followed by (41.81 cm) obtained by applying 150 kg/ha which was statistically on par. The lowest leaf length (31.23 cm) was recorded by applying no fertilizer (Table 2). Given that plant height and leaf length are influenced by the application of mineral fertilizers, this might be explained by the same factors that were previously described. The work is consistent with the findings of Getachew and Temesgen [22], noted that plots treated with 105-92-17 kg/ha N-P-S and 131.25-115-21.25 kg/ha N-P-S had obtained the longest leaf length was 24.51 cm, and 22.52 cm, respectively. Similarly, Betewulign and Solomon [25] reported that the highest leaf length (49.26 cm) was recorded by 100 kg/ha Nitrogen and 130 kg/ha P_2_O_5._

#### Yield, yield components and quality

##### Bulb length (cm)

Garlic bulb length was significantly (P < 0.01) influenced by the main factor of chicken manure and NPS fertilizer rates, but not by their interaction (Table 1).

The results showed that applying 10 t/ha of chicken manure resulted in the longest mean bulb length (2.98 cm), which was followed by 3.00 cm and 2.98 cm obtained by applying 5 and 15 t/ha, respectively, which were statistically non-significant. On the other hand, applying zero t/ha of chicken manure resulted in the shortest bulb length (2.52 cm) (Table 3). This could be because the nitrogen in chicken manure supported vegetative growth, which in turn supported photosynthesis that moved to storage organs and aided in the development of bulb length. In agreement with the current findings, Fikru and Fikreyohannes [26] reported the highest bulb length (3.31 cm) of garlic due to the application of 7.5 t/ha vermicompost. Contrasting the current study, Yohannes *et al.* [27] and [28] reported that the use of farmyard manure on onion did not show significant effect.

**Table 3:**
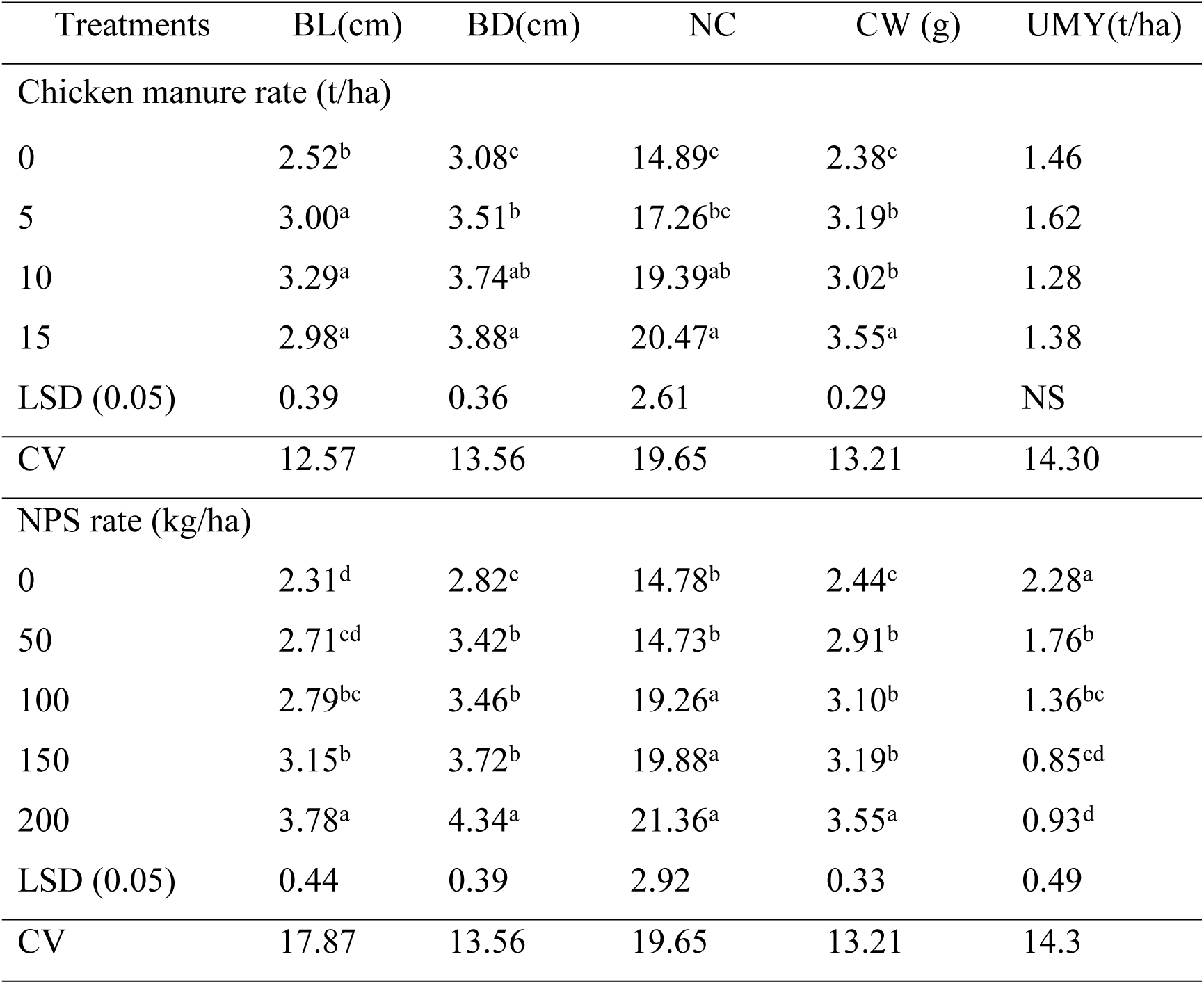
Main influence of chicken manure and NPS fertilizers rates on bulb length (BL), bulb diameter (BD), number of clove (NC), clove weight (CW) and unmarketable yield (UMY) of in 2021/2022.

The means in the column that are indicated by the same letters are not statistically different at the 5% level; CV (%) = coefficient of variation, and LSD (0.05) =least significant difference.

In terms of the main effects of NPS fertilizer, the treatment that utilized 200 kg/ha of fertilizer had the longest mean length of bulb (3.78 cm), which was followed by treatments with 150 kg/ha and 100 kg/ha of fertilizer (3.15 cm) and (2.79 cm), respectively, whereas, the fertilizer treatment with zero had got the lowest length (2.31 cm) (Table 3). This might be attributed to the availability of the applied nutrients for better growth and yield. Accordingly, Michael [24], the greatest bulb length (2.8 cm) was achieved when the soil was treated with 80 kg N/ha obtained from urea and 40 kg N substituted by cow manure. The longer garlic bulbs could be a result of increased nutritional availability, particularly nitrogen (N), which increases the length of the bulb by simulating cell enlargement and promoting plant growth [5]. Similarly, Fikru and Fikreyohannes [26] reported the highest bulb length (3.37 cm) of garlic by addition of 130 t/ha nitrogen.

##### Bulb diameter (cm)

The influence of chicken manure and NPS rates highly significantly (P < 0.01) affected bulb diameter of garlic (Table 1). Addition of treatments at the rate of 15 t/ha chicken manure resulted in the widest bulb diameter (3.88 cm) followed by a bulb diameter of 3.74 cm in a plot treated with 10 t/ha chicken manure. Both were statistically similar, whereas the smallest (3.08 cm) was recorded by zero or control treatment (Table 3). This could be the result of a sufficient supply of nutrients, which encouraged the bulb expansion. Other possibilities could include increased carbohydrate buildup, which would expand the bulb diameter. Consistent with the findings of the current experiment, Yohannes *et al*. [27], the application of FYM at a rate of 45 t/ha produced the maximum mean bulb diameter (5.99 cm). Additionally, it was noted by Mousa and Mohamed [29] that using chicken manures greatly boosted the onion bulb’s diameter.

The bulb diameter increases gradually with increasing the rates of NPS fertilizer. The plots that received 200 kg/ha showed the biggest bulb diameter (4.34 cm), whereas, the control had the lowest (2.82 cm) (Table 3). This may be explained by the nutrients N, P, and S that were present in sufficient amounts for growth, meristematic activity, and enzymatic activity, which increased bulb diameter. In consistent with the result obtained in this investigation, the findings of Assefa *et al.* [30] and [31], reported a rise in garlic bulb diameters with the administration of sulfur-containing fertilizer. Moreover, the sulfur and nitrogen in NPS fertilizer promote the production of chlorophyll and enzymatic activities, both of which lead to larger garlic bulbs [32].

##### Number of cloves

The number of cloves per bulb was significantly impacted (P < 0.01) by the main effects of chicken manure and NPS fertilizer but their interaction was not statistically significant (Table 1). Plots treated with 15 t/ha chicken manure produced the highest number of cloves per bulb (20.47), with statistically no difference (19.39) in the number of cloves obtained by 10 t/ha. The control (0 t/ha) of chicken manure yielded the lowest value (14.89) (Table 3). This could be because more chicken manure means more nutrients are released into the soil, which encourages more rapid growth and expansion of the bulb diameter and, consequently, more cloves. In agreement with the finding, Sitaula *et al.* [12], reported the highest number of cloves (24.69) per bulb in garlic by treatment supplied with 100 % poultry manure. Somen and Hitesh [17] reported that the maximum cloves number per bulb (12.33) was confirmed by 20 t/ha. Furthermore, it was demonstrated by Paczka *et al.* [33] that the addition of 50% vermicompost had the greatest positive impact on diameter, average bulb weight, number of cloves per bulb, and average clove weight.

A highly significant difference (P<0.01) in the number of cloves per bulb was also observed using NPS fertilizer. The treatment that applied 200 kg/ha of NPS fertilizer rate produced the highest number of cloves, 21.36, followed by 19.88 and 19.26 cloves, respectively, obtained by treatments of 150 and 100 kg/ha that were statistically similar. On the other hand, 50 t/ha produced the lowest number (14.73) of cloves per bulb (Table 3). This could be because the crop’s growth and productivity are dependent on receiving the ideal nutrients. The present results are consistent with those of Asefa [18], who stated that the plot receiving 157.98-137.97-25.34 kg/ha N-P-S fertilizer had the highest clove number (8.65).

##### Clove weight (g)

The results of the analysis of variance showed that the effects of NPS fertilizer and chicken manure on the weight of garlic cloves were highly significant (P < 0.01), but their interaction was non-significant (Table 1). The treatment application of 15 t/ha chicken manure produced the maximum clove weight of garlic (3.55g), followed by clove weight (3.02g) in a plot treated at a rate of 10 t/ha. Plots that were not fertilized produced the lowest weight of cloves (2.38g) (Table 3). This could be because manures enhance the fertility, structure, and nutrient availability of the soil for plants. According to Ewais *et al.* [34], improved plant nutrition accelerated photosynthesis and plant tissue cell proliferation, which is consistent with the findings of the current study. The product growth and bulb weight both show this. The outcome was consistent with the findings of Yoldas et al. [35], who observed that the parcels receiving the largest amounts of onion bulb weight were those where 60 t/ha of chicken manure had been sprayed. The treatment of 10 t/ha chicken manure to the garlic resulted in a larger weight (90.986 g) of cloves, as reported by Zakari *et al.* [36].

Garlic clove weight increased to 3.55 g when 200 kg/ha NPS was applied to the crop; the lowest weight (2.44 g) resulted with no fertilizer treatment. This could be as a result of the highest levels of sulfur, phosphorus, and nitrogen that aided in the metabolism and produced nucleic acids, phospholipids, coenzymes, and chlorophyll, all of which increase the bulb weight of garlic plants [32]. Diriba et al. [37] found a similar thing, stating that the NPS fertilizers had a major impact on the average weight of the cloves. The study also supported the findings of Fikre et al. [21], who found that the average bulb weight of onion plants across all kinds grew gradually as the rate of blended NPSB fertilizer was raised. Diriba *et al.* [37] also reported that NPS fertilizers significantly affected the mean clove weight.

#### Unmarketable bulb yield (t/ha)

The current investigation revealed that the rate of NPS fertilizer main effects had a significantly (p < 0.05) impact on the unmarketable bulb yield. The maximum unmarketable bulb yield (2.28 t/ha) was achieved by zero fertilizer application whereas the lowest 0.85 t/ha and 0.93 t/ha were recorded by the application of NPS fertilizer of 150 kg/ha and 200 kg/ha, respectively which were statistically similar (Table 3). The higher level of NPS fertilizer application indicated about a 168 % reduction in unmarketable yield as compared to zero fertilizer which has significant value as it is high-value cash crop. Similar finding was reported by Befekadu *et al.* [9]. They indicated a 57.7 % yield advantage when treated with 128 kg/ha N and 92 kg/ha P_2_O_5_ in combination with 2.5 t/ha vermicompost in comparison with zero or control.

##### Marketable bulb yield (t/ha)

According to current investigation, marketable bulb yield was highly significantly (*P < 0.01*) influenced by main effects of chicken manure and NPS fertilizer rates and their interaction (Table 1). The maximum marketable bulb yield (19.52 t/ha) and (18.53 t/ha) was obtained at the interaction of 15 t/ha chicken manure with 150 NPS kg/ha and 200 kg/ha NPS, respectively, which were statistically identical. But the least marketable bulb yield (1.90 t/ha) was recorded by control, and it was statistically on par with zero level of NPS fertilizer with 5-tone chicken manure and 50 kg/ha NPS with zero chicken manure (Table 4). The result further revealed a range value of 17.62 t/ha marketable bulb yield between maximum and minimum. Thus, the yield advantage of 927 % was observed with the higher rates of fertilizer effects as compared to zero fertilizer applications. This may be explained by the fact that fertilizers containing sulfur, such as NPS, can change the pH of the soil and enhance the availability of micronutrients, which raises the yield of garlic bulbs. Furthermore, increased fertilizer levels led to increased plant growth in terms of height, leaf length, and number of leaves, all of which increased the amount produced in garlic trough photosynthetic activity. Because sufficient nutrients lead to larger bulbs, which account for marketable yield, this raises the yield of marketable bulbs. The recent result was consistent with those of Bewuket *et al.* [9], who found that the marketable yield of garlic was greatly boosted by the combined application of vermicompost with appropriate levels of phosphate and nitrogen. Additionally, Melkamu *et al.* [38] reported that the highest marketable and total tuber yield of potatoes was achieved by combined application of 13.5 t/ha FYM and 245.1 kg/ha NPS fertilizer. Similarly, the use of zinc and boron along with N, P, K, and S in the presence of chicken manure surely enhanced garlic yield [39]. Furthermore, <u>Shege</u> *et al.* [47] in their finding reported that the highest marketable bulb yield (17.42 t/ha) was achieved by the higher doses of NPS (140:122.6:22.6) combinations in garlic plant.

**Table 4:** Chicken manure and NPS fertilizers levels influenced garlic marketable bulb yield in 2021/2022.

| NPS rate<br>(kg/ha) | Chicken manure rate (t/ha) |  |  |  |  |  |  |  |
| --- | --- | --- | --- | --- | --- | --- | --- | --- |
|  | Mean marketable bulb yield (t/ha) |  |  |  | Mean total bulb yield (t/ha) |  |  |  |
|  | 0 | 5 | 10 | 15 | 0 | 5 | 10 | 15 |
| 0 | 1.90<sup>k</sup> | 2.63<sup>jk</sup> | 3.83<sup>ij</sup> | 5.00<sup>i</sup> | 3.89<sup>k</sup> | 5.06<sup>jk</sup> | 5.95<sup>ij</sup> | 7.58<sup>jih</sup> |
| 50 | 3.22<sup>jk</sup> | 4.57<sup>i</sup> | 6.42<sup>h</sup> | 8.23<sup>g</sup> | 5.42<sup>jk</sup> | 6.58<sup>hij</sup> | 7.86<sup>fgh</sup> | 9.64<sup>e</sup> |
| 100 | 8.13<sup>g</sup> | 10.83<sup>f</sup> | 14.13<sup>cd</sup> | 17.83<sup>b</sup> | 9.75<sup>e</sup> | 12.25<sup>d</sup> | 15.41<sup>bc</sup> | 18.98<sup>a</sup> |
| 150 | 8.63<sup>g</sup> | 13.68<sup>de</sup> | 17.87<sup>b</sup> | 19.52<sup>a</sup> | 9.23<sup>efg</sup> | 14.76<sup>bc</sup> | 18.85<sup>a</sup> | 20.27<sup>a</sup> |
| 200 | 8.37<sup>g</sup> | 12.67<sup>e</sup> | 15.39<sup>c</sup> | 18.53<sup>ab</sup> | 9.31<sup>ef</sup> | 13.85<sup>cd</sup> | 15.97<sup>b</sup> | 19.55<sup>a</sup> |
| LSD (0.05) |  |  | 1.35 |  |  |  | 1.67 |  |
| CV (%) |  |  | 8.00 |  |  |  | 8.21 |  |

Means having same letter (s) either in column or row are not significantly different at 5% level, CV (%) = Coefficient of variation, LSD<sub>(0.05)</sub> = Least Significant Difference at 5% probability.

##### Total bulb yield (t/ha)

The ANOVA result revealed that the main effects of chicken manure and NPS fertilizer rates and their interaction had a highly significant (*P < 0.01*) effect on total bulb yield (Table 1). The highest total bulb yields of 20.27 t/ha, 19.55 t/ha, and 18.98 t/ha were obtained by the interaction of 15 t/ha chicken manure with 150 NPS kg/ha, 15 t/ha chicken manure with 200 kg/ha NPS, 15 t/ha chicken manure with 100 kg/ha NPS, in the given order which were statistically on par whereas, numerically the lowest (3.89 t/ha) was obtained from the control (Table 4). The investigation revealed that there was a 421 % yield advantage due to the interaction effects of fertilizers as compared to control. This may be due to the high amount of nutrients supplied from both NPS fertilizer and chicken manure. This was in agreement with the work of Driba [40] who reported the application of integrated chicken manure and inorganic fertilizers significantly increased the growth, yields and qualities of garlic. Similarly, Rizk *et al.,* [41] found the highest yield of onion attained by the application of integrated fertilizers rather than using mineral fertilizers only. Also the maximum yield (27.8 t/ha) of garlic was achieved by the addition of 50 kg/ha N with 10 t/ha manure [42] Similarly, Yadav *et al.* [1] also documented maximum bulb yield when 50 % recommended dose (120:50:100 kg/ha) of NPK applied with 12 t/ha farm yard manure.

##### Total soluble solids (°Brix)

According to the investigation, it was found that the main impacts of chicken manure and NPS fertilizer rate was statistically significant (P < 0.01) but their interaction effect had non-significant influence on the total soluble solid (Table 1). The treatment applied with 15 t/ha of chicken manure had the greatest mean total soluble solids (22.47 °Brix) followed by treatments with 10 t/ha and 5 t/ha of chicken manure that produced 20.40 °Brix and 18.73 °Brix, respectively, whereas, the lowest result (17.7 °Brix) was attained by zero/ control (Table 5). Total soluble solid was increased as increasing the level of chicken manure on the garlic plant. It might be because of increasing nutrient amount of chicken manure that contributed to more accumulation of dry matter content and reserved substances in the bulbs. Similarly, Fikru and Fikreyohannes, [26] recorded 6.7 % increase in TSS by the application of 7.5 t/ha vermicompost in garlic as compared to the control treatment. This is also supported by the report of Alemu *et al.* [5], who found more TSS in garlic due to the addition of vermicompost in contrast with the control.

**Table 5:** The influence of chicken manure and NPS fertilizers rate on total soluble solid and dry matter content of garlic in 2021/2022.

| Treatments | TSS (°Brix) | DM (%) |
| --- | --- | --- |
| Chicken manure rate (t/ha) |  |  |
| 0 | 17.7<sup>c</sup> | 26.85<sup>c</sup> |
| 5 | 18.73<sup>bc</sup> | 29.10<sup>bc</sup> |
| 10 | 20.40<sup>ab</sup> | 31.79<sup>b</sup> |
| 15 | 22.47<sup>a</sup> | 35.31<sup>a</sup> |
| LSD<sub>(0.05)</sub> | 2.39 | 3.32 |
| NPS rate (kg/ha) |  |  |
| 0 | 16.50<sup>c</sup> | 22.11<sup>c</sup> |
| 50 | 18.42<sup>bc</sup> | 29.99<sup>b</sup> |
| 100 | 20.75<sup>ab</sup> | 33.31<sup>ab</sup> |
| 150 | 21.92<sup>a</sup> | 32.16<sup>b</sup> |
| 200 | 21.25<sup>a</sup> | 36.25<sup>a</sup> |
| LSD<sub>(0.05)</sub> | 2.68 | 3.72 |
| CV | 16.42 | 14.62 |

Values with the same letters in the column are not significantly different at 5% level, CV (%) = coefficient of variation, LSD<sub>(0.05)</sub> = Least Significant Difference at 5% probability level.

The NPS fertilizers rate significantly (P < 0.01) influenced the total soluble solid of garlic. The maximum total soluble solid (21.92 °Brix) was obtained by 150 kg/ha NPS whereas the smallest total soluble solid (16.50 °Brix) was attained by zero fertilizer (Table 5). This might be attributed to the low level of dry matter concentration with zero fertilizer application. This result was consistent with the research conducted by Fismes *et al.* [43], which showed that the addition of sulfur fertilizer resulted in bulbs with a greater TSS content. Garlic is known for its ability to absorb larger amounts of accessible nutrients, particularly sulfur nutrients. In a similar vein, Diriba *et al*. [37] reported that the total soluble solids (TSS) of garlic bulbs were significantly affected by the application of N, P, and S fertilizers. In contrast, the lowest TSS result (14.20°Brix) was produced by bulbs without any fertilizer application. Furthermore, Bloem *et al.* [44] found that during major growth until the start of ripening, the content of aliins in garlic was dramatically raised by high sulfur quantity treatment combined with low nitrogen fertilization. On the other hand, elevated nitrogen levels had a negative impact.

#### Bulb dry matter weight (%)

The current study revealed that NPS fertilizer rates and chicken manure main effect had significantly (*P < 0.01*) influenced bulb dry matter content. However, their interaction was non-significant (Table 1). Relatively higher percent of dry matter yield (35.31 %) was observed application of (15 t/ha) of chicken manure, followed by (31.7%) and (29.10%) that received 10 t/ha and 5 t/ha of chicken manure, respectively. Conversely, the lowest (26.85 %) was recorded by zero application of chicken manure (Table 5). A higher level of chicken manure has shown a yield advantage of 31.5 % increment as compared to zero application. This result was also supported by Fikru and Fikreyohannes [26] who reported the significant influence of vermicompost on bulb dry matter content of garlic. They recorded 32.71 % dry matter yield by the application of 7.5 t/ha as compared to 27.01 % obtained by zero vermicompost. In contrast to this study, Ayed [45] revealed that chicken manure had no significant difference in dry matter weight of onion.

The addition of 200 kg/ha NPS fertilizer gave the highest bulb dry matter percent (36.25%), but the lowest bulb dry matter (22.11%) was recorded by zero fertilizer treatment. Due to NPS fertilizer treatment, the crop has shown a range of 14.14 with an advantage of 63.95 % (Table 5). This could be the impact of nutrients that are essential for the accumulation of dry matter. Similarly, Diriba *et al.,* [37] found the effects of N, P, and S fertilizers on bulb quality, namely dry matter percentage, total soluble solids, pungency, bulb protein content, and weight loss in storage organ, and concentration of important nutrients in the bulb tissues. The dry matter found in this study were similar to the result of Sardia and Timra, [46] who reported the maximum shoot and root dry mass of garlic when NPSK application increased over the control.

#### Economic return

The results showed that the combined effect of NPS mineral fertilizer and chicken manure at different amounts and types of combination for garlic production was significantly influenced by cost of input that vary, gross and net returns of garlic bulb. Higher net benefits (524, 224.8 ETB/ ha) were obtained by the combined application of 15 t/ha chicken manure and 150 kg/ha NPS mineral fertilizer. The result of economic analysis further revealed the highest marginal rate of return of (10,159.1 %) by combined application of 15 t/ha chicken manure and 150 kg/ha NPS fertilizer followed by a marginal rate of return 9, 658.5 % (421,343.2 ETB/ ha net benefit) by the combination of 5 t/ha chicken manure and 150 kg/ha NPS (Table 6).

**Table 6:** Summary of economic analysis for response of chicken manure and NPS fertilizers rate of garlic in Lemo woreda, Hadiya zone, Southern Ethiopia, 2021/2022.

| CM rate<br>( t ha<sup>-1</sup>) | NPS rate<br>( kg ha<sup>-1</sup>) | MY<br>(kg ha<sup>-1</sup>) | AMY<br>(kg ha<sup>-1</sup>) | TVC<br>(Birr ha<sup>-1</sup>) | GFB<br>(Birr ha<sup>-1</sup>) | NB<br>(Birr ha<sup>-1</sup>) | MRR % |
| --- | --- | --- | --- | --- | --- | --- | --- |
| 0 | 0 | 1,900 | 1710 | 2394 | 68400 | 66006 | - |
| 0 | 50 | 3,220 | 2898 | 4957.2 | 115920 | 110962.8 | 1753.9 |
| 0 | 100 | 8,130 | 7317 | 12043.8 | 292680 | 280636.2 | 2394.3 |
| 0 | 150 | 8,630 | 7767 | 13573.8 | 310680 | 297106.2 | 1076.5 |
| 0 | 200 | 8,370 | 7533 | 14146.2 | 301320 | 287173.8 | D |
| 5 | 0 | 2,630 | 2367 | 54513.8 | 94680 | 40166.2 | D |
| 5 | 50 | 4,570 | 4113 | 57858.2 | 164520 | 106661.8 | 1,988.3 |
| 5 | 100 | 10,830 | 9747 | 66645.8 | 389880 | 323234.2 | 2,464.5 |
| 5 | 200 | 12,670 | 11403 | 70764.2 | 456120 | 385,355.8 | 1,508.4 |
| 5 | 150 | 13,680 | 12312 | 71136.8 | 492480 | 421,343.2 | 9,658.5 |
| 10 | 0 | 3,830 | 3447 | 106025.8 | 137880 | 31854.2 | D |
| 10 | 50 | 6,420 | 5778 | 110189.2 | 231120 | 120,930.8 | 2,139.5 |
| 10 | 100 | 14,130 | 12717 | 120803.8 | 508680 | 387,876.2 | 2,514.9 |
| 10 | 200 | 15,390 | 13851 | 124191.4 | 554040 | 429,848.6 | 1,239.0 |
| 10 | 150 | 17,870 | 16083 | 126416.2 | 643320 | 516,903.8 | 3,912.9 |
| 15 | 0 | 5,000 | 4500 | 157500 | 180000 | 22,500 | D |
| 15 | 50 | 8,230 | 7407 | 162469.8 | 296280 | 133,810.2 | 2,239.7 |
| 15 | 100 | 17,830 | 16047 | 175465.8 | 641880 | 466,414.2 | 2,559.3 |
| 15 | 200 | 18,530 | 16677 | 178147.8 | 667080 | 488,932.2 | 839.6 |
| 15 | 150 | 19,520 | 17568 | 178495.2 | 702720 | 524,224.8* | 10,159.1* |

MY= Marketable yield, AMY= adjusted marketable yield GFB = gross field benefit; TVC = total variable costs; NB = net benefit, MRR = marginal rate of return; ETB ha<sup>-1</sup> = Ethiopian Birr ha<sup>-1</sup>; D = dominated treatments, NPS cost =18 Birr kg<sup>-1</sup>, Chicken manure = 10 Birr kg<sup>-1</sup>, Labor cost fertilizer application = 1200 Birr ha<sup>-1</sup>, Material cost= 200 ETB t<sup>-1</sup>, transportation = 500 ETB t<sup>-1</sup>, Harvesting = 700 ETB t<sup>-1</sup> and market price of garlic = 40 Birr kg<sup>-1</sup> during harvesting time at farm gate price.

According to CMMYT [16] an MRR higher than 100 % is generally acceptable for the production and productivity of crops under given conditions; it is better to maximize the net benefit and MRR to its highest possible level. In this regard, although the statistics have shown non-significant differences in treatment combination of 15 t/ha chicken manure in combination with 150 and 200 kg/ha NPS fertilizers, the economic analysis revealed the higher value of using 150 kg/ha NPS fertilizer as compared to 200 kg/ha. Similar results has been reported by Getachew and Asefa [11], who indicated the highest net return of Ethiopian Birr 626,814 per hectare with the combined application of 112–37–16 kg ha^−1^ NPS and 15 t ha^−1^ cattle manure.

## Conclusion and recommendation

The current investigation results clearly revealed that the productivity of garlic responded positively and significantly to integrated applications of chicken manure and NPS mineral fertilizers. The data indicated that the response of garlic to higher levels of both organic and inorganic fertilizers was significant for growth, yield, and quality parameters investigated. The application of 15 t/ha chicken manure and 150 to 200 kg/ha NPS fertilizer gave higher values in most of growth parameters like plant height, leaf number, leaf lengths, and leaf diameter. In the case of yield components, yield, and quality parameters, the combined application of 15 t/ha and 150 kg/ha NPS was found to be optimum in most of the investigated parameters, with some exceptions. The quality parameters such as bulb dry matter and TSS were also found to be significantly higher when 15 t/ha chicken manure and about 150 NPS kg/ha fertilizer were applied. The economically most important parameter, marketable bulb yield, was found to be statistically non-significant for treatment combinations of 15 t/ha chicken manure with 150 and 200 kg/ha NPS, further economic analysis confirmed the better net income of Ethiopian Birr (currency) 524, 224.8 Birr/ha) with the highest marginal rate of return of 10,159.1 % achieved by the combination of 15 t/ha chicken manure and 150 kg/ha NPS mineral fertilizer. Therefore, it could be recommended that the combined application of 15 t/ha chicken manure with 150 kg/ha NPS fertilizers can be used for economically feasible production and productivity of garlic in the study area. Also, the usage of integrated fertilizers has additional benefits for soil properties and the environment as well.

## Author contributions

Abrham Shumbulo – corresponding author, prepared the manuscript, participated in field research; Mihret Jemal – conducted the field research, data collection and organization, Abraham Bosha – contributed in data analysis, manuscript write up and proof readings.

## Data availability

All scientifically important data are included in the body of the text.

## Acknowledgments

The authors sincerely acknowledge Wachamo University for the support in laboratory facilities, field experiments, and financial support for some running costs during field experiments. In addition, Wolaita Sodo University, Department of Horticulture deserves thanks for the cooperation in some running costs facilities.

## References

1. Yadav RN, Bairwa HL and MK Gurjar Response of garlic (*Allium sativum* L.) to organic manures and fertilizers. International Journal of Current Microbiology and Applied Sciences, 2017; 6(10):4860–4867.

2. FAO (Food and Agriculture Organizatioof United Nations) Area and production of crops by countries.”, 2018;”Available: www.faostat.fao.org.

3. CSA (Central Statistical Agency) Agricultural Sample Survey. Report on area and production of crops, 2020; Addis Ababa, Ethiopia.

4. Kilgori MJ, Magaji MD and AI Yakubu Effect of plant spacing and date of planting on yield of two garlic (*Allium sativum* L.) Cultivars in Sokoto, Nigeria, American Eurasian. Journal of Agricultural and Environmental Science, 2007; 2: 153–157.

5. Alemu D, Nigussie D and G Fikreyohannes Effects of vermicompost and inorganic NP fertilizers on growth, yield and quality of garlic (*Allium sativum* L.) in EnebseSarMidir District, Northwestern Ethiopia. *Journal of Biology*, Agriculture and Healthcare, 2016; 6(3): 22243208

6. Yusuf A, Fagbuaro SS and SO Fajemilehin Chemical composition, phytochemical and mineral profile of garlic (*Allium sativum*). Journal of Bioscience and Biotechnology Discovery, 2018; 3(5):105–109.

7. Abay A Assessment of soil fertility status of different types of soils in selected areas of Southern Ethiopia. Journal of Natural Sciences Research, 2016; 6(1):2224-3186.

8. Belay Y Integrated soil fertility management for better crop production in Ethiopia. International Journal of Soil Science, 2015; 10:1-16.

9. Befekadu T, Missiganaw W and A Endeshaw The traditional practice of farmers’ legume-cereal cropping system and the role of microbes for soil fertility improvement in North Shoa, Ethiopia. Agricultural Research and Technology, 2017; 13(4):1–6.

10. Bewuket G, Kebede W and B Ketema Effects of organic and inorganic NP fertilizers on the performance of garlic (*Allium sativum* L.) varieties at Koga, Northwestern Ethiopia. *Journal of Biology*, Agriculture and Healthcare, 2017; 7(7):2224–3208.

11. Getachew A and A Asefa Influence of integrated soil fertilization on the productivity and economic return of garlic (*Allium sativum* L.) and soil fertility in northwest Ethiopian highlands. De Gruyter Open Agriculture, 2021; 6: 714–727. 10.1515/opag-2021-0047

12. Sitaula HP, Dhakal R, Bhattarai C, Aryal A and D Bhandari1 Effects of different combinations of poultry manure and urea on growth, yield and economics of garlic (*Allium sativum* L.). Journal of Agriculture and Natural Resources, 2020; 3(1): 253–264 ISSN: 2661-6289 (Online) DOI: 10.3126/janr.v3i1.27179

13. Islah MEl-H Response of Garlic (*Allium sativum* L.) to some Sources of Organic Fertilizers under North Sinai Conditions. Resources of Journal Agriculture and Biology Science, 2010; 6(6): 928-936.

14. Ghanbarian D, Youneji S, Fallah S and A Farhadi Effect of broiler litter on physical properties, growth and yield of two cultivars of cantaloupe (Cucumismelo) International. Journal of Agricultural Biology, 2008; 10: 697–700.

15. Getachew T and Z. Asfaw Achievements in shallot and garlic research. Report No.36. Ethiopian Agricultural Research Organization, 2000; Addis Ababa. Ethiopia.

16. CIMMYT From Agronomic Data to Farmer Recommendations: An Economics Training Manual. Revised Edition. 1988; D.F. Mexico 51p.

17. Somen A and K Hitesh Effect of Some Organic Manure on Growth and Yield of Garlic in Greenhouse Condition at Cold Desert High Altitude Ladakh Region, Defence Life Science Journal, 2018; 3(2): 100–104

18. Asefa A Effect of Cattle Manure and NPS Mineral Fertilizer on the Productivity of Garlic (*Allium Sativum* L.) using irrigation in Lay Gaint District, North Western, Ethiopia. 2017;

19. Adewale OM, Adebayo OS and T A Fariyike Effect of Poultry Manure on Garlic (*Allium sativum* L) Production in Ibadan, South Western Nigeria. Continental J. Agricultural Science, 2011; 5(2): 7 – 11.

20. Indupulapati M Essential plant nutrients and their functions, Center of International de Agricultural Tropical (CIAT), 2009; Cali, Colombia, 36.p

21. Fikre K, Yohannes DB and AN Woldekirstos Effects of blended (NPSB) fertilizer rate on growth, yield and yield component of onion (*Allium cepa* L.) varieties at Jimma condition by Wolkite, Ethiopia. AGBIR, 2021; 37(4):152–157.

22. Getachew A and M Temesgen Effects of Nitrogen and NPS Fertilizer Rates on Fresh Yield of Garlic (*Allium sativum* L.) at Debre Berhan, Ethiopia, Journal of Agriculture and Crops, 2020; 6(8): 113–118, DOI: 10.32861/jac.68.113.118

23. Frank GV The plants need for and use of Nitrogen in soils. Edited by Bartaacovew W.C, Francis E.C. Amer. Soc. of Agronomy, 2000; M.C USA. pp 508 – 509

24. Michael Akuamoah-Boateng Effect of Poultry Manure and NPK Fertilizer on the Growth, Yield and Mineral Composition of Onion (*Allium cepa* L.) Grown in Acid Soil, Msc. Thesis, 2016; the University of Ghana, Legon. http://ugspace.ug.edu.gh/

25. Betewulign E and T Solomon Evaluating the Role of Nitrogen and Phosphorous on the Growth Performance of Garlic (*Allium sativum* L.). Asian Journal of Agricultural Research, 2014; 8: 211–217.

26. Fikru T and G Fikreyohannes Response of Garlic (*Allium sativum* L.) Growth and Bulb Yield to Application of Vermicompost and Mineral Nitrogen Fertilizers in Haramaya District, Eastern Ethiopia, East African Journal of Sciences, 2019; 13(2): 159–168

27. Yohannes KW, Belew D and A. Debela Effect of farmyard manure and nitrogen fertilizer rates on growth, yield and yield components of onion (*Allium cepa* L.) at Jimma, Southwest Ethiopia. – Asian Journal of Plant Sciences, 2013; 12(6-8): 228–234.

28. Mekonnen DA, Mihretu FG and K Woldetsadi Farmyard manure and intra-row spacing on yield and yield components of Adama red onion (*Allium cepa* L.) cultivar under irrigation in Gewane district, Afar Region, Ethiopia. Horti. For, 2017; 9(5): 40–48.

29. Mousa MA A and MF Mohamed Enhanced yield and quality of onion (*Allium cepa* L.) produced using organic fertilization. Assiut University Bulletin for Environmental Researches. 2009; 12(1): 9 - 19.

30. Assefa AG, Mesgina SH and YW Abrha Effect of inorganic and organic fertilizers on the growth and yield of garlic crop (*Allium sativum* L.) in Northern Ethiopia. Journal of Agricultural Science, 2015; 7: 80–86.

31. Hariyappa N Effect of potassium and sulfur on growth, yield and quality parameters of onion (*Allium cepa* L.) (M.Sc.thesis), 2003; University of Agricultural Sciences, Dharwad, India.

32. Shafie F and E Gamaily Effect of organic manure, sulfur and microelements on growth, bulb yield, storability and chemical composition of onion plants. Minufiya Journal of Agricultural Research, 2002; 27: 407– 424.

33. Paczka G, Mazur-Pączka A, Garczyńska M, Kostecka J and KR Butt Garlic (*Allium sativum* L.) Cultivation Using Vermicompost Amended Soil as an Aspect of Sustainable Plant Production. Sustainability, 2021, 13, 13557

34. Ewais MA, Mahmoud AA and AA Khalil Effect of nitrogen fertigation in comparison with soil application on onion production in sandy soils. – Alex J Agriculture Res, 2010; 55(3): 75–83.

35. Yoldas F, Ceylan S and N. Mordogan Effect of Chicken Manure on Yield and Yield Criteria of Onion (*Allium cepa* L.) As Second Crop. Applied Ecology and Environmental Research, 2019; 17(5):12639–12647.

36. Zakari S, Miko M and BS Aliyu Effect of Different Types and Levels of Organic Manures on Yield and Yield Components of Garlic (*Allium sativum* L.) At Kadawa, Kano, Nigeria, Bayero. Journal of Pure and Applied Sciences, 2014; 7(1): 121 – 126.

37. Diriba S, Nigussie D, Kebede W, Getachew T and JJ Sharma The effect of Nitrogen, Phosphrus and Sulfur Fertilizers on growth, yield and economic returns on garlic (Allium sativum L.) Conditions. Journal of Applied Science Research. 2015; 10(5):383–392.42.

38. Melkamu A, Minwyelet J, Tadele Y and A Masho Integrated application of compound NPS fertilizer and farmyard manure for economical production of irrigated potato (*Solanum tuberosum* L.) in highlands of Ethiopia, Cogent Food & Agriculture, 2020; 6(1): 1724385.

39. Nasreen MN, Yousuf AN, Mamun M, Brahma S and M M Haquc Response of Garlic to Zinc, Boron and Poultry Manure Application. Bangladesh J. Agril. Res. 2009; 34(2): 239–245. DOI: 10.3329/bjar.v34i2.5795

40. Driba S Review of Management Strategies of Constraints in Garlic (*Allium sativum* L.) Production. Journal of Agricultural Sciences, 2016; 11(3):186-207.

41. Rizk FA, Shaheen AM, El-Samad EH and T T El-Labban Response of onion plants to organic fertilizer and foliar spraying of some micro-nutrients under sandy soil. Journal of Applied Sciences Research. 2014; 10(5):383–392. http://www.aensiweb.com/jasr.html

42. Tadila G and Nigusie D Effect of manure and nitrogen rates on growth and yield of garlic (*Allium sativum* L.) at Haramaya, Eastern Ethiopia, Journal of Horticulture and Forestry, 2018; 10(9):135–142. 10.5897/JHF2018.0543

43. Fismes J, Vong PC, Gucker A, and E. Frossard Influence of sulfur on apparent N-use efficiency, yield and quality of oilseed rape (*Brassica napus* L.) grown on a calcareous soil. Eur. J. Agron. 2000; 12:127–141.

44. Bloem E, Haneklaus S and E Schnug Storage Life of Field-Grown Garlic Bulbs (*Allium sativum* L.) as Influenced by Nitrogen and Sulfur Fertilization. J. Agric. Food Chem. 2011; 59(9): 4442–4447.

45. Ayed A Effect of Chicken manure, Sheep manure and Inorganic fertilizer on yield and nutrient uptake by onion, Pakistan Journal of Biology science, 2002;5(3):266-268.

46. Sardia K and E Timra Responses of Garlic (*Allium sativum* L.) to Varying Fertilization Levels and Nutrient Ratios. J. Commun. Soil Sci. Plant Anal, 2005; 36(4 & 6):673- 679.

47. Yayeh, S. G., Alemayehu, M., Haileslassie, A., Dessalegn, Y., & Tejada Moral, M. Economic and agronomic optimum rates of NPS fertilizer for irrigated garlic (*Allium sativum L*) production in the highlands of Ethiopia. Cogent Food & Agriculture, 2017; 3(1):1–10. 10.1080/23311932.2017.1333666

